# Single-cell atlas of human oral mucosa reveals a stromal-neutrophil axis regulating tissue immunity in health and inflammatory disease

**DOI:** 10.1101/2021.04.06.438702

**Authors:** DW Williams, T Greenwell- Wild, L Brenchley, N Dutzan, A Overmiller, AP Sawaya, S Webb, D Martin, G Hajishengallis, K Divaris, M Morasso, M Haniffa, NM Moutsopoulos

## Abstract

The oral mucosa remains an understudied barrier tissue rich in exposure to antigens, commensals and pathogens. Moreover, it is the tissue where one of the most prevalent human microbe-triggered inflammatory diseases, periodontitis, occurs. To understand this complex environment at the cellular level, we assemble herein a human single-cell transcriptome atlas of oral mucosal tissues in health and periodontitis. Our work reveals transcriptional diversity of stromal and immune cell populations, predicts intercellular communication and uncovers an altered immune responsiveness of stromal cells participating in tissue homeostasis and disease at the gingival mucosa. In health, we define unique populations of *CXCL1,2,8-*expressing epithelial cells and fibroblasts mediating immune homeostasis primarily through the recruitment of neutrophils. In disease, we further observe stromal, particularly fibroblast hyper-responsiveness linked to recruitment of leukocytes and neutrophil populations. Ultimately, a stromal-neutrophil axis emerges as a key regulator of mucosal immunity. Pursuant to these findings, most Mendelian forms of periodontitis were shown to be linked to genetic mutations in neutrophil and select fibroblast-expressed genes. Moreover, we document previously unappreciated expression of known pattern- and damage-recognition receptors on stromal cell populations in the setting of periodontitis, suggesting avenues for triggering stromal responses. This comprehensive atlas offers an important reference for in-depth understanding of oral mucosal homeostasis and inflammation and reveals unique stromal–immune interactions implicated in tissue immunity.

## Introduction

The oral mucosa is a site of first encounters^1^. Life-sustaining substances such as food, water, and air pass through the oral cavity, carrying along allergens, microbes, and food particles as they enter the gastrointestinal tract or respiratory system. In addition to constant exposure to microbes, allergens and other antigens, tissues in the oral cavity must withstand frequent microdamage from masticatory forces. Despite – or perhaps because of – this tumultuous environment, the oral barrier displays minimal infection at mucosal surfaces and no overt inflammation at steady state. Moreover, the oral mucosa is remarkably efficient at wound healing^2^, as most injuries heal without infection or scar formation. Yet, the mechanisms by which oral mucosal tissues interpret and respond to environmental cues are still largely unappreciated, which renders the study of this remarkably exposed and resilient tissue of broad interest.

The oral mucosal barrier is a multilayer squamous cell epithelium. However, different areas within the oral cavity have acquired unique anatomical characteristics adapted to the functions and specialization of their specific microenvironment^3^. The majority of the oral mucosal barrier consists of a multilayer squamous epithelium with minimal keratinization (lining epithelium) which lines the inside of the cheeks (buccal mucosa) (Fig.1a, right enlarged inset), the inside of lips, the floor of the mouth and the back of the throat (soft palate), and protects important organs, such as the salivary glands and large vessels and nerves that supply the craniofacial structures. The rest of the oral mucosa is specialized to perform specific functions. As such, the hard palate, tongue dorsum and external area of gingiva are fully keratinized to withstand the constant mechanical forces of mastication (masticatory epithelia) and the tongue also contains taste buds to mediate taste perception. A particularly unique area of the oral barrier is the gingiva, the tooth-associated mucosa (Fig. 1a left enlarged inset). The external area of the gingiva is formed by masticatory epithelium, while the internal side (gingival crevice) is a non-keratinized squamous epithelium, which becomes progressively thin (down to 3-4 layers at its base). This particularly thin mucosa is also constantly exposed to the complex microbial biofilm at the tooth surface^4^, and is potentially the most vulnerable site for microbial translocation within the oral cavity^3^.

**Figure 1:**
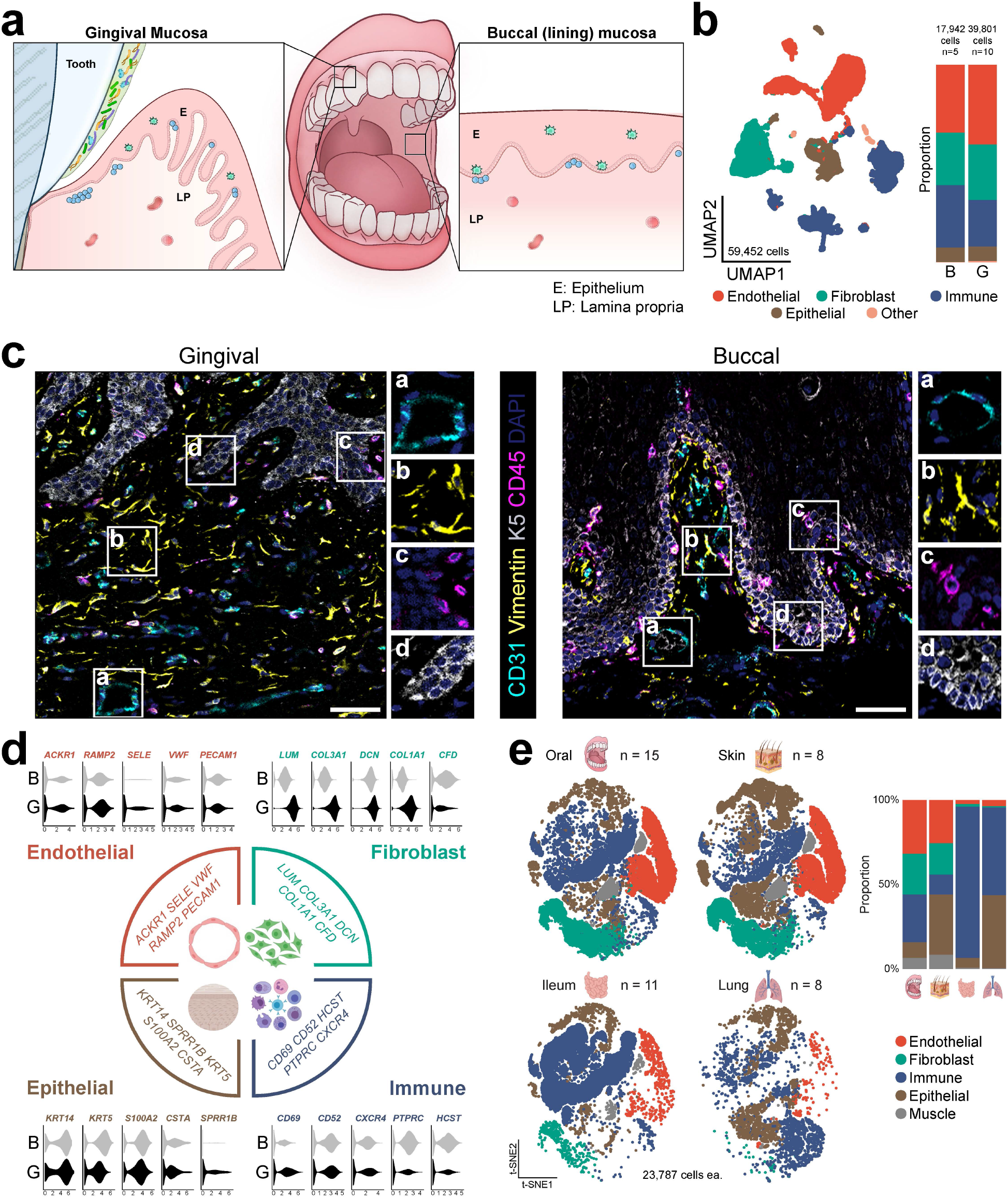
Major cell types in healthy adult oral mucosal tissues. **a**. Schematic of the oral mucosal tissues biopsied for this study with their respective locations indicated within the oral cavity (E-epithelium, LP-lamina propria). **b**. UMAP representation of major cell types identified by scRNA Seq (n=15, 59,452 cells, left) and bar graph of relative cell proportions by tissue type (B-buccal, G-gingival) (right). **c**. Immunofluorescence depicting major cell types in healthy gingival and buccal mucosa tissue. Scale bar: 41μm. Stains indicated and color coded. **d**. Violin plots of selected markers for major cell populations identified by scRNA Seq and relative expression per tissue type. X-axis represents z-scored average expression. **e**. t-SNE (left) and proportion plots (right) showing representation of major cell types in oral, skin, ileum, and lung barrier tissues. Demographics of datasets utilized in this analysis shown in Supplemental Table 6

In fact, a major distinction in oral epithelia can be made between buccal and gingival mucosae. Indeed, the commensal microbiome communities are particularly rich, diverse and complex at the tooth interface compared to microbial communities found in all other oral sites^5,6^, suggesting that the gingival mucosa may be specifically suited to interact with microbes. Furthermore, disease susceptibilities within the oral mucosa are most distinct between the lining and gingival mucosa. As such, ulcerative disorders and infections are most commonly present at the lining mucosa, while bullous dermatological diseases are encountered at the external gingiva^7^. Finally, the gingiva is also the location for one of the most common inflammatory human diseases, periodontitis.

Periodontitis is a highly prevalent human disease, affecting in its moderate-to-severe forms approximately 8% of the general population^8^. In periodontitis, a dysbiotic microbiome on the tooth surface is thought to trigger exaggerated inflammation within the tooth associated mucosa (gingiva) in susceptible individuals^9,10^. This dysregulated inflammatory response leads to destruction of tooth-supporting structures, including mucosal tissues and supporting bone. To date the pathogenic mechanisms involved in periodontitis are incompletely understood.

Herein, we have compiled a human cell atlas of oral mucosal tissues in health and periodontal disease. Our work entails a comprehensive characterization of cell populations and their transcriptional signatures in the setting of clinical health and inflammatory disease and aims to uncover cell types and functions associated with promoting homeostasis and disease susceptibility at this important barrier. Indeed, through this atlas we identify unique populations of stromal cells promoting neutrophil migration in health and expanding in disease. Furthermore, evaluation of cellular expression of genes linked to periodontal -disease susceptibility further reveals stromal and immune cell (particularly neutrophil) involvement in disease susceptibility. Collectively, our work provides a cellular census of distinct oral mucosal tissues in health and disease and suggests unique stromal-immune responsiveness mediating mucosal homeostasis which becomes dysregulated in mucosal inflammatory disease.

## Results

### Single-cell sequencing of ∼60,000 cells identifies 4 major tissue compartments in healthy oral mucosa

To create a comprehensive single-cell transcriptome atlas of the human oral mucosa, we obtained biopsies of the buccal and gingival mucosa from healthy volunteer s meticulously screened for systemic and oral health (Fig. 1a, Supplemental table 1). Biopsies were processed immediately into a single-cell suspension by mild enzymatic digestion and mechanical separation, followed by library preparation and sequencing using the 10x Genomics Chromium Droplet platform. After quality control to remove low-quality cells and cells expressing high mitochondrial gene signatures, our dataset included 39,801 cells from the gingival mucosa (n=10) and 17,942 cells from the buccal mucosa (n=5) (Supplemental Fig. 1a-c). Our analysis revealed 4 major cell compartments in oral mucosa comprised of epithelial, endothelial, fibroblast and immune cells, which were present in both gingiva and buccal mucosa of healthy adults (Fig. 1b). Histological imaging confirmed the presence of the aforementioned 4 major cellular compartments within oral mucosal tissues and provided insight into the tissue architecture of oral mucosal tissues (Fig. 1c).

Analysis of the transcriptomic signatures for the major cell types documented differential expression of cell-defining genes for the 4 major cell compartments, which were generally conserved across mucosal sites. In fact endothelial (*ACKR1, RAMP2, SELE, VWF, PECAM1*), fibroblast (*LUM, COL3A1, DCN, COL1A1, CFD*), immune (*CD69, CD52, CXCR4, PTPRC, HCST*), and epithelial (*KRT14, KRT5, S100A2, CTSA, SPRR1B*) cell genes were consistent with classical, well-established markers for each cell population (Fig. 1d, Supplemental Fig. 1d).

Finally, interrogation of published data sets from healthy barrier tissues across the human body (oral, skin^11^, ileum^12^, lung^13^) demonstrated that these 4 major populations are also the main cellular compartments in all human barrier tissues evaluated (Fig. 1e). However, the relative representation of each major cell population differed between tissue types. Representation of these 4 main building blocks of barrier tissues was most comparable between the oral mucosa and skin barriers, although the oral mucosa contained an expanded immune compartment while the skin harbored a considerably larger epithelial compartment (Fig. 1e).

### Diverse stromal and epithelial subpopulations with distinct immune functionality are present in oral mucosal tissues

Sub-clustering within the 4 main cell types demonstrates great diversity in stromal, epithelial and immune cell populations in human oral mucosal tissues, with a total of 34 transcriptionally distinct clusters (Supplemental Fig. 2a).

Within the endothelial cell compartment, we defined 4 health-associated (H) venular endothelial cell subsets (H.VEC), a population of vascular smooth muscle cells (H.SMC) and a population of lymphatic endothelial cells (H.LEC), all of which were represented in both buccal and gingival mucosa (Fig. 2a, Supplemental Table 2). H.SMC and H.LEC clusters express genes related to muscle contraction (such as *MYL9, ACTA2*) and LEC fate commitment (*PROX1*) (Fig. 2b, Supplemental Fig. 2b). Venular endothelial cell subpopulations expressed genes consistent with distinct functions. H.VEC 1.1, 1.3, and 1.4 were all linked to immune functions. H.VEC 1.1 expressed a gene signature consistent with antigen presentation (*HLADRA, CD74*), while 1.3 and 1.4 appeared to promote immune and inflammatory responses (such as response to LPS and IFN signaling) and expressed genes relevant to neutrophil recruitment and IFN signaling (*CSF3, CXCL2, MX1)* (Fig. 2b, Supplemental Fig. 2b). H.VEC 1.2 cells displayed a gene signature consistent with active pathways for endothelium development (such as *NOTCH4, SOX18*) (Fig. 2b, Supplemental Fig. 2b).

**Figure 2:**
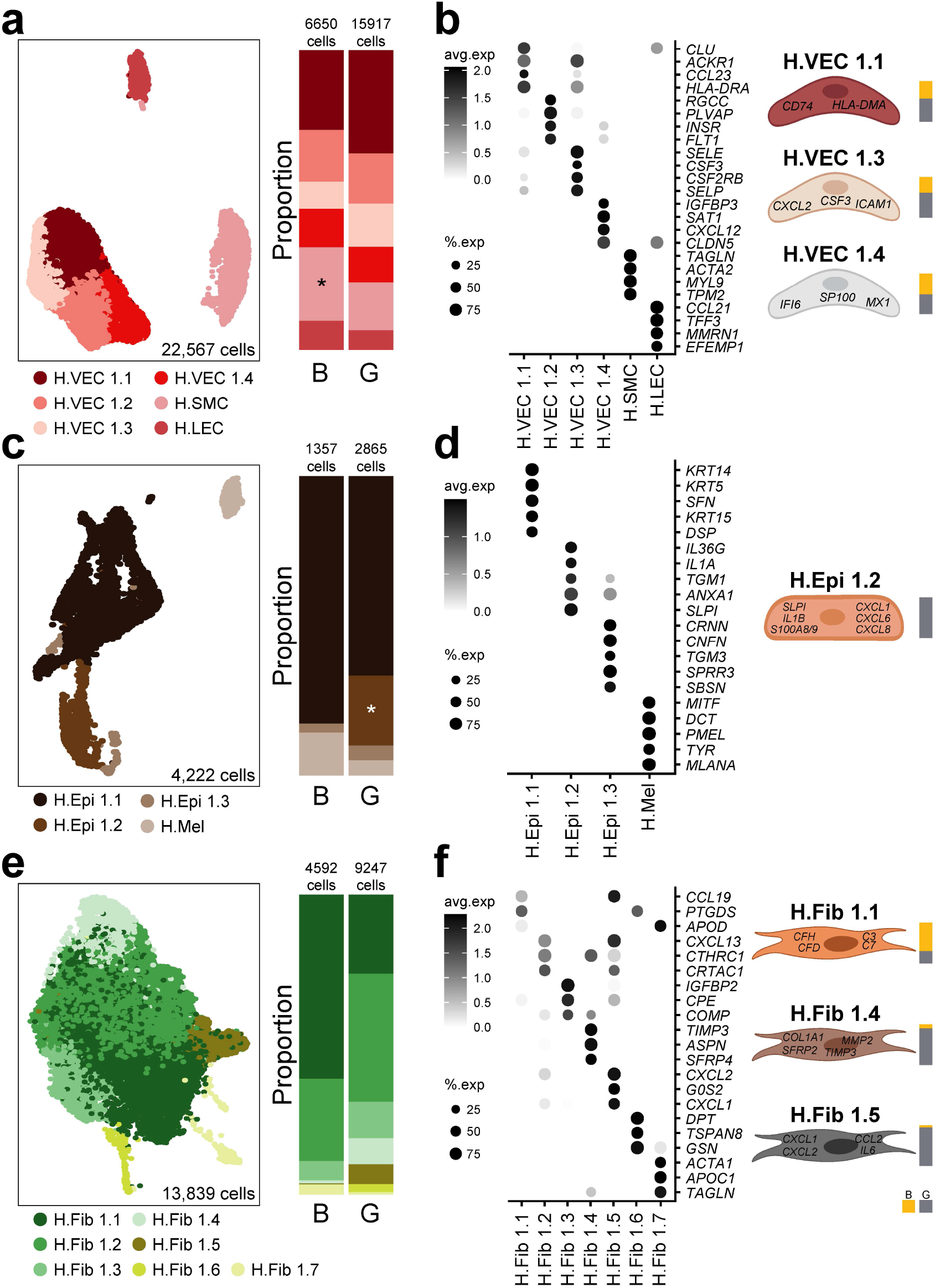
Stromal and epithelial subpopulations in healthy oral mucosa. **a**,**c**,**e**. UMAP (left) and proportion plots (right) of endothelial (22,567 cells), epithelial (4,222 cells) and fibroblast (13,839 cells) populations, respectively, present in gingival (G) and buccal (B) mucosa. Refer to methods for statistical tests used. *p < 0.05. **b**,**d**,**f**. Dot plots depicting the expression of cluster-defining genes and percentage of cells expressing each gene for endothelial, epithelial, and fibroblast populations, respectively. Expression values are Z-scored averages. To the left of each dot plot is a cartoon illustrating the subtypes that express genes associated with immune activation and recruitment. Proportion plots to the right of each illustration depict the contribution of each tissue type to each cell subtype.

Within the epithelial compartment, we identified 4 epithelial cell subtypes in healthy oral mucosa; three subtypes of keratinocytes (H.Epi 1.1,1.2,1.3) and a melanocyte population (H.Mel) (Fig. 2c,d, Supplemental Table 2). H.Mel was the most distal epithelial cluster by UMAP (Fig. 2c) and clearly represented melanocytes based on significant expression of pigmentation genes (*PMEL, MLANA*) (Fig. 2d, Supplemental Fig. 2c). Based on gene expression, H.Epi 1.1 cells were further categorized as basal layer/differentiating keratinocytes with expression of *KRT5, KRT14* and *KRT15*, while H.Epi 1.3 expressed genes involved in keratinization and cornification (*SPRR3, CRNN, CNFN*) typically encountered in the outmost layers of the epithelium (Fig. 2d, Supplemental Fig. 2c). Whereas H.Epi 1.1 and 1.3 were present in both buccal and gingival mucosa, H.Epi 1.2 was only found in the gingiva and featured a gene expression profile consistent with inflammatory responses. In fact, top expressed genes included antimicrobial and inflammatory factors such as *IL1b, SLPI*, and *S100A8/9*, and a top pathway was leukocyte recruitment, which reflected primarily recruitment of neutrophils as evident by the expression of *CXCL1, CXCL6, CXCL8* (Fig. 2d, Supplemental Fig. 2c).

Fibroblasts separated into 7 cell clusters which displayed gene signatures consistent with distinct inferred functions (Fig. 2e, Supplemental Table 2). Fibroblast clusters H.Fib 1.2, 1.3, and 1.4 displayed a gene signature corresponding to the prototypic functions of the cell type, that is, extracellular matrix organization, skeletal morphogenesis and tissue remodeling, while clusters H.Fib 1.1, 1.5, and 1.6 expressed gene signatures consistent with inflammatory responses, such as complement activation (*C3, CFD*), response to LPS (*CXCL1, 2, 8*) and development-associated BMP signaling pathways (*WNT5A*) (Fig. 2f, Supplemental Fig. 2d). H.Fib 1.7 was characterized as a myofibroblast-like population based on expression of related genes (*TAGLN, ACTA1)* and was present primarily in healthy buccal mucosa. Interestingly, “immune-like” fibroblast clusters H.Fib 1.5 (transcriptome consistent with innate immune responses and recruitment of neutrophils) and H.Fib 1.4 (tissue remodeling, *MMP2, TIMP3*) were expanded in the gingiva compared to the buccal mucosa (Fig. 2f).

Overall, our data reveal a complex landscape of epithelial and stromal cells are present in oral mucosal tissues that have diverse inferred functions. Importantly, expansion of epithelial and stromal cells with inflammatory signatures in the gingiva suggested that innate immune responses, particularly related to recruitment of neutrophils and active tissue turn over, are potentially mediated by the stromal compartment at the gingival mucosa

### T and myeloid cells are dominant in healthy oral mucosa

Within the immune cell compartment of the healthy oral mucosa we found a rich and diverse population of immune cells. Generally, immune cells divided into 5 major clusters of T/NK, B/plasma, and granulocyte/myeloid cells, which were defined accordingly by classic cell-specific marker gene expression (Fig. 3a, Supplemental Table 3). Immune cells were annotated using SingleR and manual annotation, which identified 14 unique cell types that were confirmed based on gene expression for marker genes (Fig. 3a-d). In both buccal and gingival mucosa, the majority of immune cells were T cells, which were subdivided into αβ CD4^+^, T_H_17, mucosal-associated invariant T (MAIT), αβ CD8^+^, γδ T, T_reg_, and NK/T cells. The second largest overall population was myeloid cells, which included myeloid dendritic cells (mDC), macrophages and neutrophils. We also observed a population of mast cells, and small populations of plasmacytoid DC (pDC), B and plasma cells (Fig. 3a-d). Buccal mucosa harbored an increased proportion of MAIT and CD8^+^ T cells whereas proportions of T_reg_, mast, and B cells were elevated in the gingiva. Consistent with the sequencing data, flow cytometry on biopsies from an independent cohort of volunteers (n=10-14, Supplemental Table 4) revealed an overall enriched immune compartment in the gingiva with significantly increased proportion and number of CD45^+^ cells in gingival biopsies (Figure 3e). Flow cytometry also confirmed the presence of major immune cell populations in healthy oral mucosa including CD45^+^CD3^+^ T cells, CD45^+^CD19^+^ B cells, CD45^+^HLADR^+^ antigen presenting cells (APC), HLADR^-^CD15^+^SSC^high^ and HLADR^-^ CD117^+^SSC^high^ granulocytes (Fig. 3f-h, Supplemental Fig. 3a). Increased inflammatory cell numbers in the gingival mucosa were primarily attributed to the increased presence of granulocytes (mast cells and neutrophils) and B cells in gingiva compared to buccal mucosa, consistent with sequencing results.

**Figure 3:**
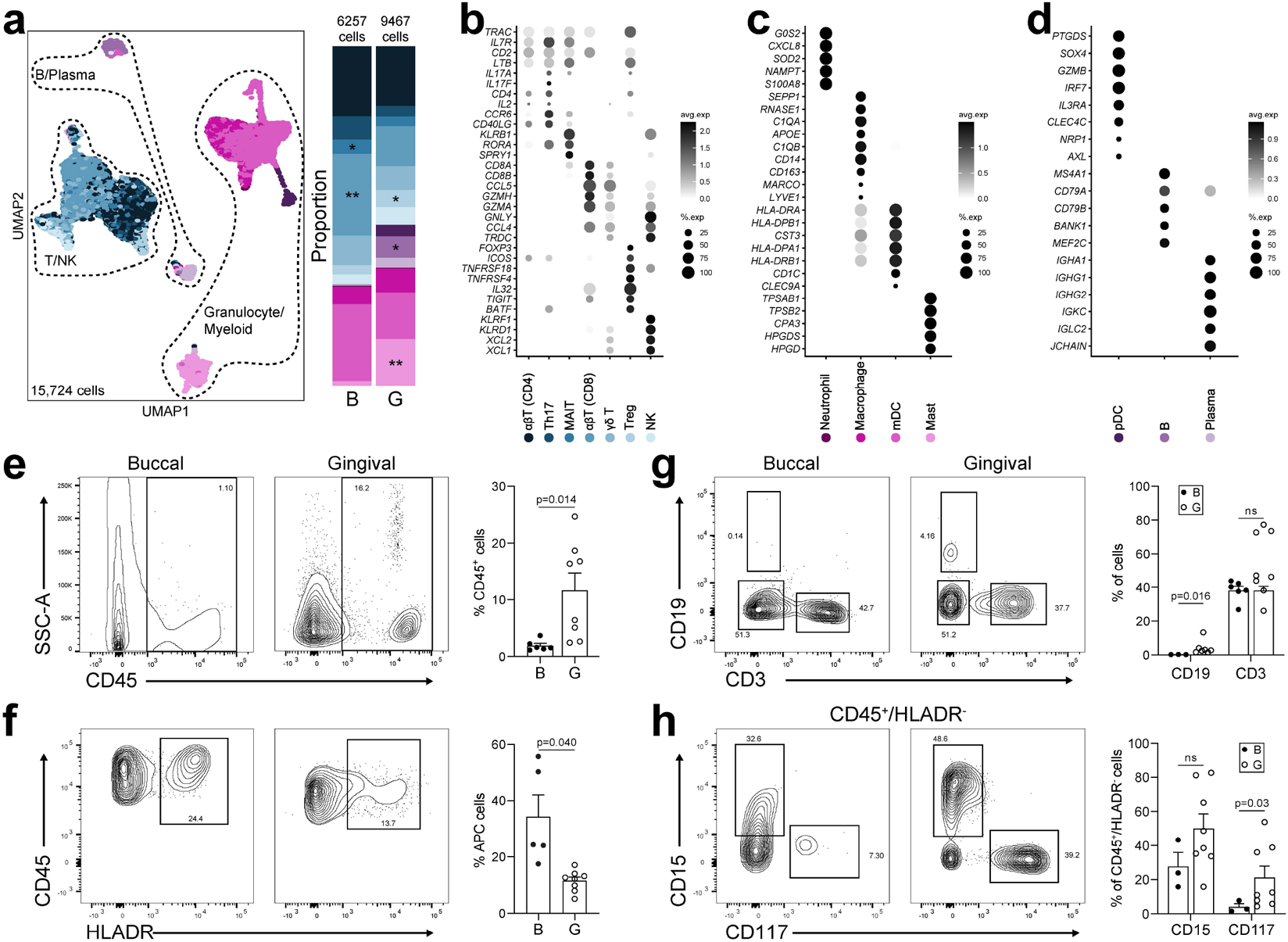
Immune populations in healthy oral mucosa. **a**. UMAP (left) and proportion plots (right) of immune cell populations in oral mucosa (15,724 cells; gingival (G), buccal (B)). Immune cells were annotated using a combination of reference based (SingleR) and manual annotation. Refer to methods for statistical tests used. *p < 0.05, **p<0.01. **b-d**. Dot plots depicting expression of cell type marker genes. **e-h**. Representative flow cytometry scatter plots from buccal (n=3-8) and gingival (n=8) biopsies. Cells were gated from Single/Live and stained with CD45 **(e)** and CD45/HLADR **(f)**, CD45^+^ cells were further stained with CD19/CD3 **(g)** and HLADR^-^SSC^high^CD15/CD117 **(h)**. Bar graphs **(e-h)** demonstrate % expression for each marker indicated. Refer to methods for statistical tests used. p values indicated on each graph.

Finally, we interrogated immune cell populations in published databases of healthy barrier tissues across the human body (oral, ileum, skin, lung). We found representation of the immune cell subsets present in the oral mucosa across other barriers, yet with varying proportions. Interestingly, we documented a significantly higher proportion of neutrophils at the oral mucosa compared to other barriers, suggesting active innate immune responsiveness at this constantly stimulated barrier, and consistent with the unique phenotype of neutrophil-recruiting stromal cells at the oral barrier (Supplemental Fig. 3b).

Altogether, this data illustrates the complex cellular constituency of the oral mucosa at steady state and reveals unique and expanded cell populations at the gingival mucosa specialized towards neutrophil recruitment and active tissue turn over. Moreover, cell-type specific transcriptomes reflect the unique anatomical characteristics of this mucosal site, which is subject to microbial stimulation and constant mechanical triggering.

### Immune cell infiltration of mucosal tissues is the hallmark of periodontitis

Host responses at the gingiva become exaggerated and destructive in the setting of the common human inflammatory disease, periodontitis. Therefore, we next aimed to characterize cell populations and their transcriptomes at the single-cell level in the setting of periodontitis. For these studies, gingival mucosal biopsies from healthy adults were compared to biopsies from patients with untreated, severe periodontitis via scRNA seq and complementary histology and flow cytometry studies (Fig. 4a). We document profound infiltration of inflammatory cells in the mucosal tissues of periodontitis with these independent approaches. H&E and immunofluorescence staining in healthy and diseased tissues revealed a profound infiltration of mucosal tissues with CD45^+^ leukocytes in the setting of disease (Fig. 4b, c). Significantly increased inflammatory cell representation was also confirmed by flow cytometry in an independent group of healthy and diseased individuals (n=6/group) (Fig. 4d). Finally, integration of health with diseased scRNA seq datasets (n=15, ∼54,000 cells) revealed 31 transcriptionally distinct clusters and a profound increase in immune cells within lesions of disease with a corresponding reduction in stromal cells, reflecting mucosal tissue destruction in periodontitis (Fig. 4e, Supplemental Fig. 4a).

**Figure 4:**
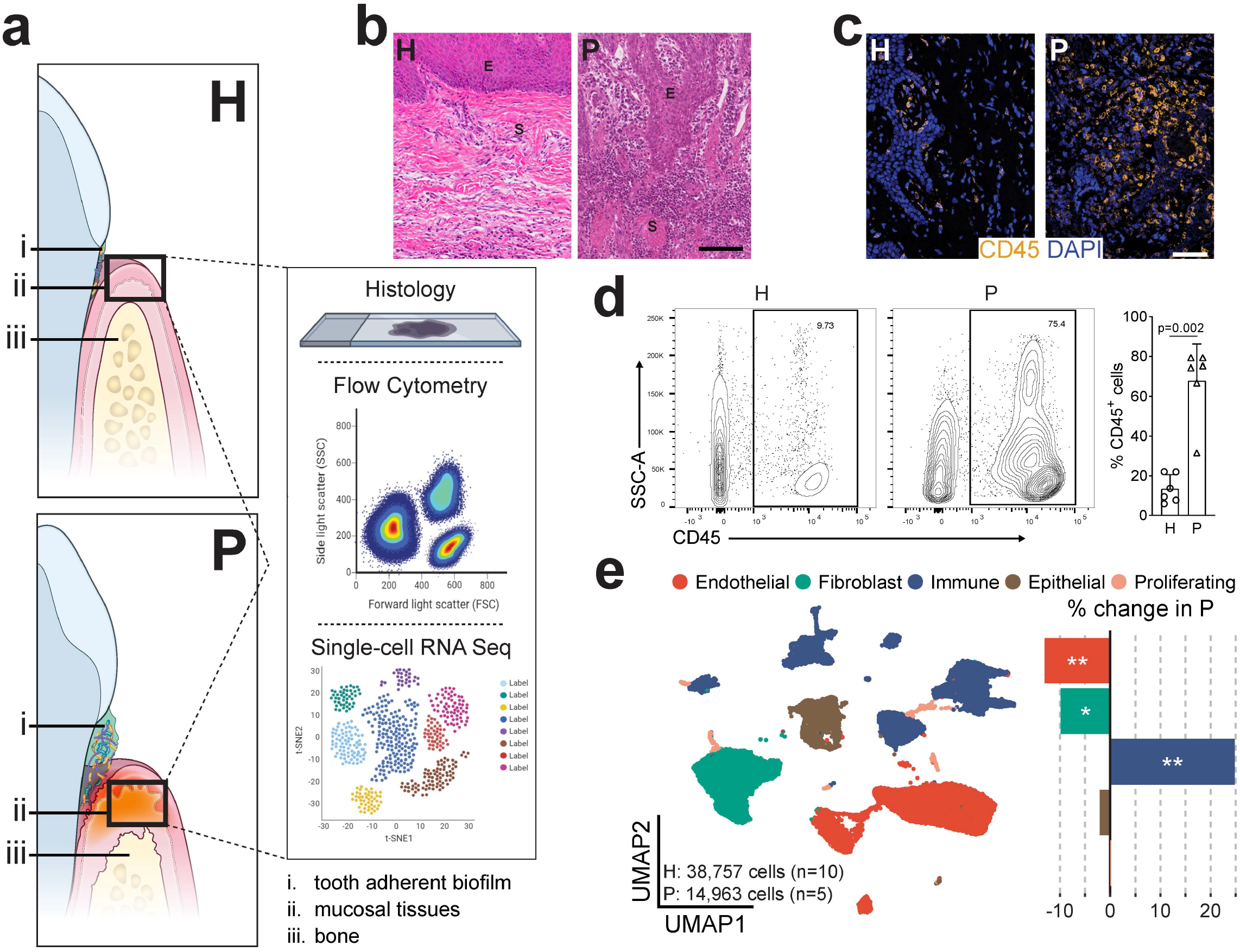
Periodontal disease is an inflammatory disease at the gingival oral mucosa. **a**. Illustration of the gingival mucosa in health (H, top) and periodontal disease (P, bottom). i) tooth adherent biofilm, ii) mucosal tissues, iii) bone. A schematic of methods for tissue analysis is present (right). **b**. H&E staining of representative gingival health (left) and periodontitis (right) sections. E-epithelium, S-stroma. Scale bar: 100μm. **c**. Immunofluorescence staining for CD45^+^ lymphocytes. Scale bar: 50μm. **d**. Representative flow cytometry scatter plots (left). Cells were gated from Single/Live and stained with CD45, graph indicates CD45 % expression (one dot per patient, n=6/group. Refer to methods for statistical tests used, p value indicated). **e**. UMAP representation of major cell populations in health and periodontal disease (left; H: 38,757 cells, n=10; P: 14,963 cells, n=5). Graph demonstrates % change in cell proportions with disease (right). Refer to methods for statistical tests used. *p < 0.05, **p<0.01.

Investigation of the immune cell types in health and disease demonstrated the presence of the same immune cell subsets previously seen in health, albeit with varying proportions in disease. Similar to health, T cells remained the major immune cell population with representation of αβ CD4^+^, T_H_17, MAIT, αβ CD8^+^, γδ T, T_reg_, and NK/T cells in disease (Fig. 5a). The next largest populations were B and plasma cells, which were significantly expanded in periodontitis compared to health (Fig. 5a, b). Finally, myeloid/granulocyte cell populations were also present in disease, with expansion of neutrophils (p=0.1429) (Fig. 5a, b). Focusing on the cell types that expand in disease, we further define gene signatures for plasma cells and neutrophils in periodontitis. A majority of plasma cells expressed IgG and either κ or λ light chains while a small but distinct population expressed IgA (Fig. 5c). Neutrophils did not subdivide into further populations based on top expressed genes but appear to uniformly express classical neutrophil markers such as *CSF3R, CXCR2 and FCGR3B* (Fig. 5d).

**Figure 5:**
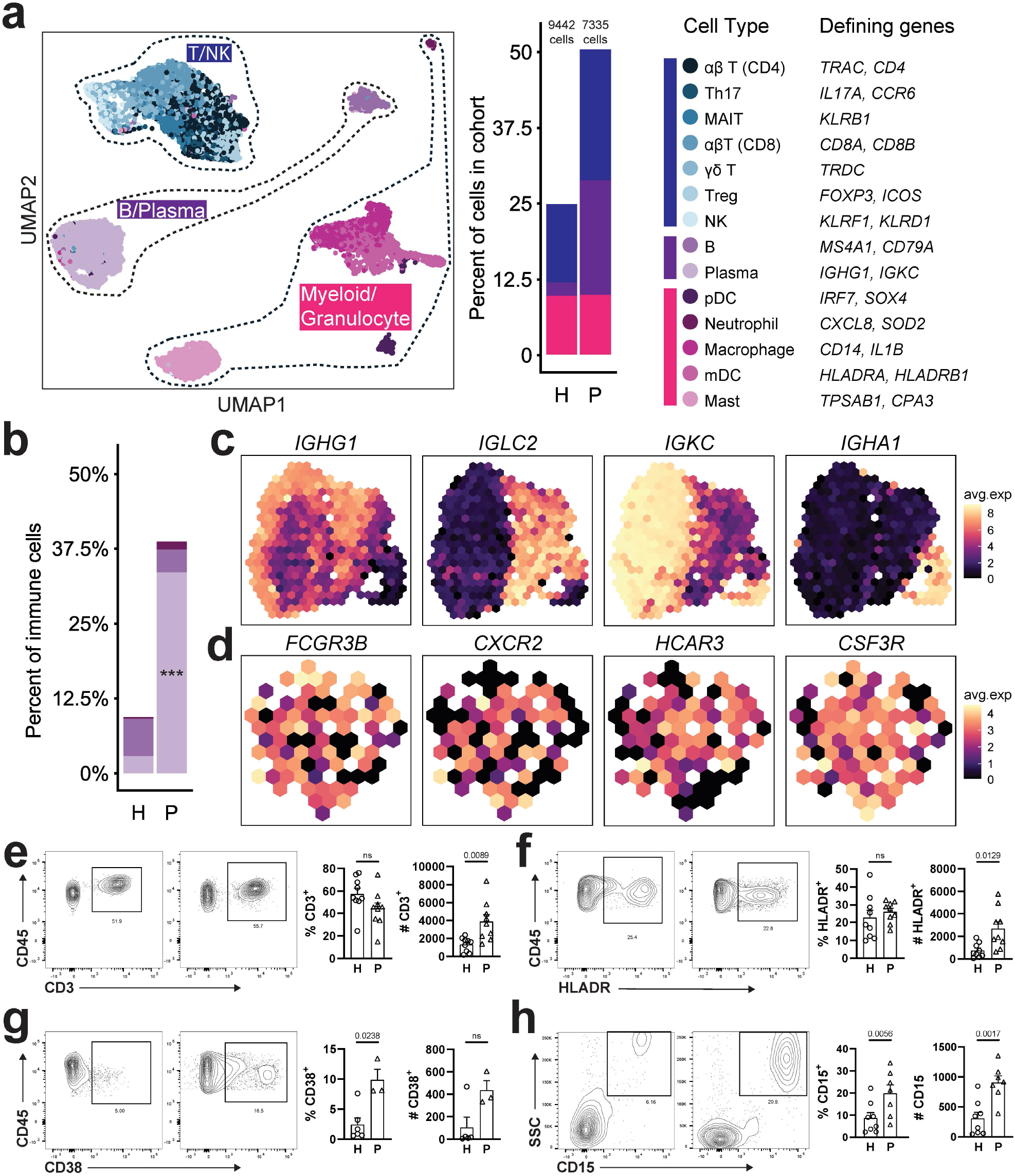
Immune cell subpopulations in periodontal disease. **a**. UMAP (left) showing immune cell subpopulations divided spatially into three main categories (T/NK, B/Plasma and Myeloid/Granulocyte). Proportion plot (middle). Colors indicate the cell type annotated using SingleR and manual annotation and validated by cell-type specific gene expression (right). **b**. Bar graph depicts proportion of B, plasma cells and neutrophils in health (H) and periodontitis (P). Refer to methods for statistical tests used. ***p<0.001. **c-d**. Z-scored average expression of plasma cell-**(c)** and neutrophil-**(d)** specific markers visualized in low-dimensional space with schex. Each area containing cells on the UMAP was divided into hexagonal areas and cells within each area were averaged. **e-h**. Representative flow cytometry scatter plots from an independent health and periodontitis cohort. Cells were gated from Single/Live cells and stained with CD45/CD3 **(e)**, CD45/HLADR **(f)**, CD45/CD38 **(g)**, CD45/CD15 **(h)**. Bar graphs demonstrate % expression (one dot per individual, n=3-9/group). Refer to methods for statistical tests used. P values are indicated on each graph.

Flow cytometry in an independent patient cohort (n=3-9 individuals/group) demonstrated significantly increased numbers of CD3^+^ T cells, CD38^+^ plasma cells, HLADR^+^ APC, and HLADR^-^SSC^high^CD15^+^ neutrophils in periodontitis (Fig. 5e-h). However, only plasma cells and neutrophils were increased in proportion with disease, consistent with data from single-cell sequencing. Of interest, while scRNA seq revealed an increase in neutrophil numbers in periodontitis, it did not capture the magnitude of neutrophil infiltration seen by flow and histology (Figure 5h, Supplemental Fig. 4b,c), reflecting a relative limitation of this method in capturing neutrophil transcriptomes^14^.

### Stromal and epithelial cells display transcriptional signatures of inflammation in periodontitis

We next investigated changes in the stromal and epithelial compartments in disease. Overall, both histological and scRNA seq analysis showed that epithelial and stromal cell proportions decreased in disease (Fig. 4b,e). Investigation of cell subsets in the combined health and periodontitis gingival database uncover novel clustering of stromal/epithelial compartments (now indicated as P, Fig. 6) with shifts in select cell subsets in disease. Periodontitis led to a significant decrease in P.VEC 1.1 (p=0.0244), a population of endothelial cells expressing antigen-presenting molecules, and an increase in P.VEC 1.2 (p=0.0280), endothelial cells characterized by expression of trans-endothelial transport gene *PLVAP* (Fig. 6a, Supplemental Fig. 5a). Notable disease-associated changes included an increase in P.Epi 1.1,the basal layer like/regenerating epithelial cells, and an increase in P.Fib 1.1,a subset of fibroblasts with expression inflammatory chemokines such as *CXCL2* and *CXCL13* (Fig. 6b,c, Supplemental Fig. 5b,c). Overall, the gene expression profile of epithelial and stromal cells shifted to an inflammatory state in disease. Specifically, endothelial cells upregulated pathways related to cell adhesion, and lymphocyte adhesion (Fig. 6d). Epithelial cells and fibroblasts upregulated pathways related to antimicrobial responses (response to LPS and response to molecule of bacterial origin) and cytokine biosynthesis, respectively (Fig. 6e,f). Of interest, genes associated with antimicrobial processes upregulated in both fibroblasts and epithelial cells in disease reflected active neutrophil recruitment (*CXCL1, CXCL2, CXCL8*) (Supplemental Fig. 5c, Supplemental Table 5). This data suggests unique wiring of the epithelial and stromal compartment in periodontitis particularly geared towards neutrophil recruitment.

**Figure 6:**
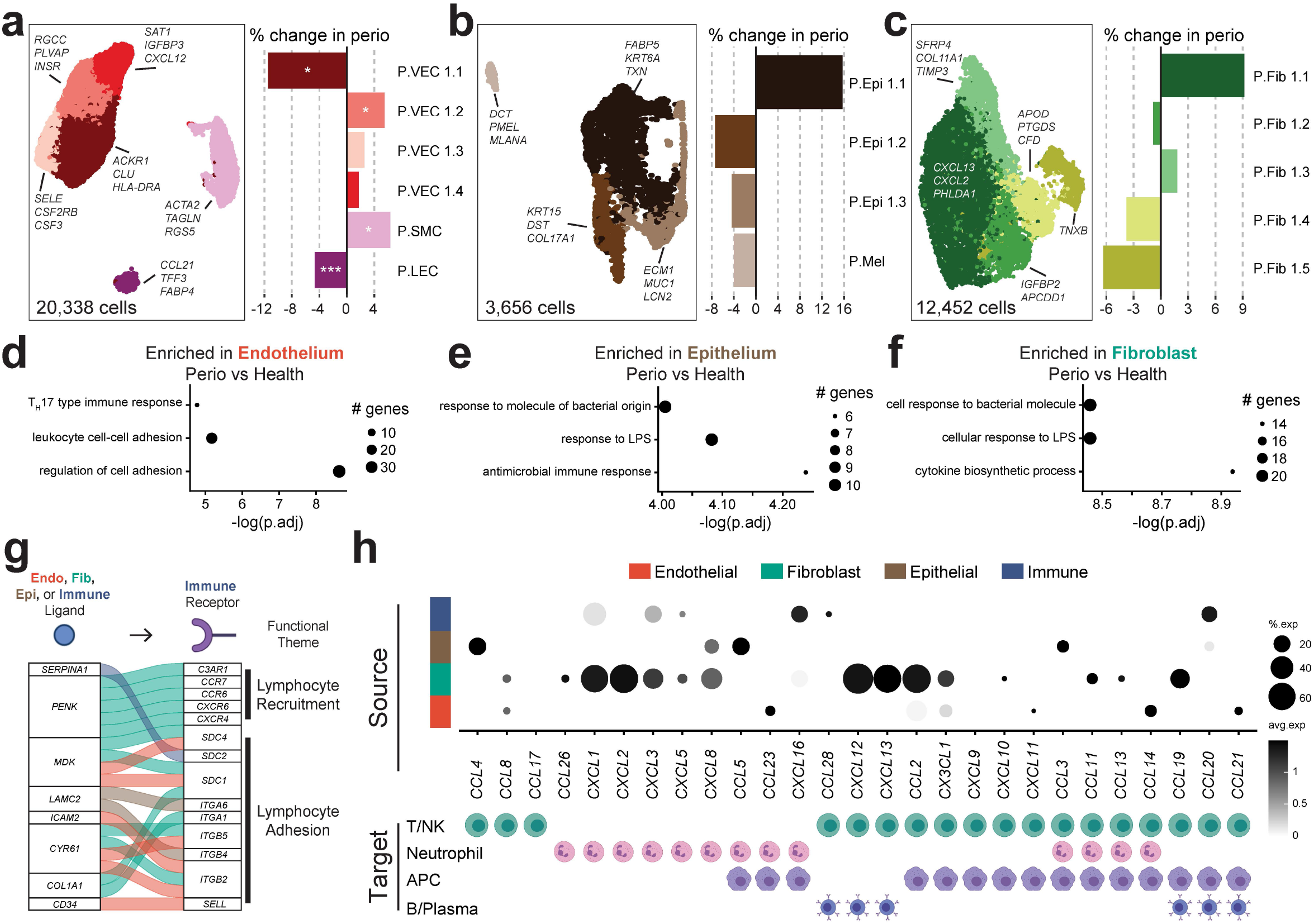
Stromal cell populations and stromal-immune interactome in periodontitis. **a**,**b**,**c**. UMAP representation (left) and proportion plots (right) for endothelial (20,338 cells), epithelial (3,653 cells) and fibroblast (12,452 cells) subpopulations, respectively, in the combined health (H) and periodontitis (P) dataset. Gene labels indicate cluster-defining genes. Refer to methods for statistical tests used. *p < 0.05. **d**,**e**,**f**. Dot plots of pathways enriched in periodontal disease per major cell population. GO terms truncated for brevity, original terms and gene list can be found in Supplemental Table 5. **g**. Alluvial plot showing selected ligand-receptor pairs significantly over-represented in periodontal disease determined by NicheNetR. For this analysis only interactions of immune cells (receptors) with all other cell types are considered. **h**. Chemokine/chemokine receptor interactions in health and periodontal disease. Dot plot indicating expression of chemokines in major cell types (top) and the immune cell(s) in the scRNA Seq dataset that express cognate receptors (bottom). Refer to Supplemental Fig. 7 for expression data of chemokine receptors.

### Stromal-immune cell interactions are potential drivers of periodontal inflammation

To further investigate potential stromal- and epithelial-immune interactions in the context of periodontitis we used NicheNet to infer cell-cell interactions with a prior model of ligand-target regulatory potential. We identified the top predicted immune receptor-ligand interactions based on gene expression in our scRNA seq dataset that were differentially regulated in disease. In periodontitis, there was extensive inferred communication between stromal and immune cells (Supplemental Fig. 6). Of interest and consistent with pathways upregulated in disease, endothelial cells appeared to promote adhesion of immune cells, while fibroblasts displayed a potential towards recruitment of inflammatory cells (Fig. 6g). Overall, based on gene expression signatures and predicted interactions, our data indicates an active role for stromal cells in the recruitment of immune cells to the site of disease.

To further explore the overall potential of stromal cells to recruit immune cells, we specifically interrogated expression of chemokine and chemokine receptor pairs in our dataset (Fig. 6h, Supplemental Fig. 7). Our data showed broad expression of chemokine ligands by all cell types in disease. In line with pathway analysis (Fig. 6f), fibroblasts were particularly transcriptionally active in the production of chemokines. Fibroblasts expressed a broad array of chemokine ligands exclusive in their potential to recruit neutrophils (*CXCL1, 2, 5, 8*) as well as chemokines with the potential of recruiting several types of leukocytes (*e*.*g. CXCL12, CXCL13, CCL19*) (Fig. 6h). Collectively, this data suggests that stromal cells utilize intercellular signaling to drive immune cell recruitment and tissue transmigration in periodontitis.

### Single-cell expression of periodontal disease-related genes suggests roles for both the immune and stromal compartment in disease susceptibility

To begin to understand cell contribution to disease pathogenesis, we next investigated cellular expression of genes linked to periodontitis susceptibility. Select Mendelian genetic diseases are clearly linked to periodontal disease susceptibility^15-17^ and recent GWAS studies have uncovered single-nucleotide polymorphisms associated with periodontal disease^18,19^. However, the cellular expression patterns and role of the relevant genetic determinants in periodontitis remains unclear. Our scRNA seq atlas uniquely enabled us to globally assess the expression of 15 extant periodontal disease genes within the gingival mucosa. Mendelian disease genes associated with periodontitis were almost exclusively expressed in the immune cell compartment with the exception of *C1S* and *C1R* (expressed exclusively in fibroblasts) indicating that immune and potentially fibroblast cell dysfunction is a critical driver of monogenic periodontitis. Neutrophil-related defects (*HAX1, LYST, ITGB2, FPR1*) are particularly linked to susceptibility for periodontitis in genetic syndromes. In contrast, disease-associated genes identified by GWAS^20-26^ were more broadly expressed, including by stromal, epithelial and immune cells, thereby suggesting the involvement of multiple pathways in the common forms of periodontal disease (Fig. 7a). Our exploration of disease associated genes at the tissue level highlight potentially diverse cell populations involved in common forms of disease, with unique roles for neutrophils and fibroblast populations in the pathogenesis of Mendelian forms of periodontitis.

**Figure 7:**
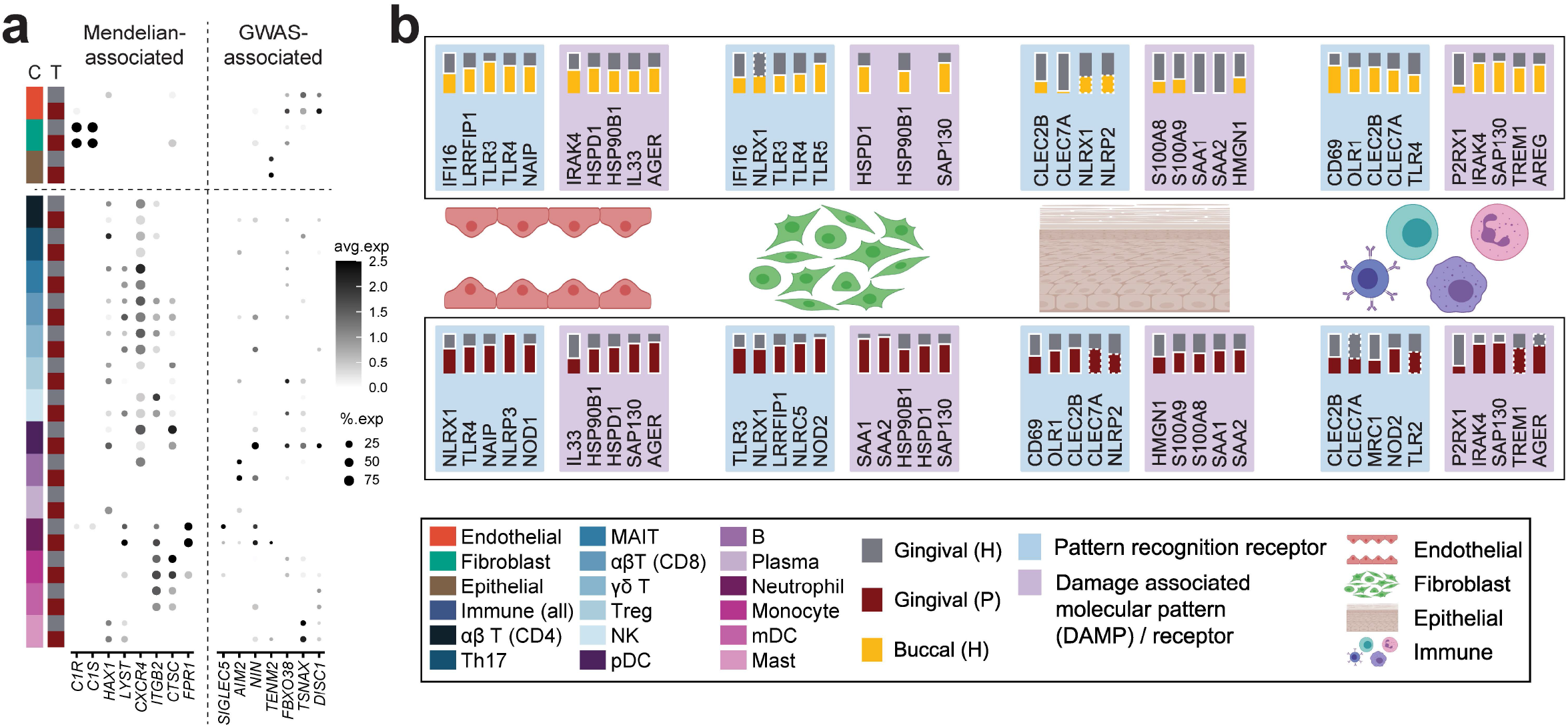
Genes associated with periodontal disease susceptibility, microbe and damage sensing in the oral mucosa. **a**. Dot plot depicting the Z-score average expression of genes and percentage of expressing cells for genes associated with Medellian forms of periodontitis (left) and genes identified by GWAS to be associated with periodontitis (right). **b**. Summary of select pattern recognition receptors and damage-associated molecular pattern and associated receptors expression in buccal and gingival healthy mucosa (top) and expression in gingival health and periodontitis (bottom). Expression is shown by major tissue type (Endothelial, Fibroblast, Epithelial, Immune). A solid white border indicates statistical significance and the tissue that had higher expression. A dotted border indicates non-significance and the tissue that had a trending increase in expression. Significance determined by non-parametric Wilcoxon rank sum test (Supplemental Table 7, 8).

### Microbiome and damage sensing may underlie stromal hyperresponsiveness in periodontitis

Finally, to investigate triggering of stromal-immune cell interactions linked to periodontitis we utilize the atlas to investigate expression of microbial and cell-damage receptors and sensors within our oral cell database in health and disease. As such we explore the cellular distribution of pattern recognition receptors, including Toll like receptors (TLR), C-type lectin receptors (CLR), nucleotide-binding oligomerization domain-like receptors (NLR), RIG-I-like receptors (RLR), viral entry factors, and of damage associated molecular patterns (DAMP) and associated receptors (DAMP-R) (Fig. 7b, Supplemental Fig. 8). We document diverse expression of microbial and damage receptors in epithelial, endothelial, fibroblast and immune cells within oral tissues in health consistent with microbial and/or damage triggering of immune responsiveness in the stromal and immune compartments. Furthermore, we find significantly increased expression of NLR, CLR, TLR, DAMP and DAMP-R molecules in disease, particularly within stromal and epithelial cells suggesting increased triggering of these cellular populations in the disease setting.

Collectively, our data promote the concept that stromal cell populations in the gingival mucosa are constantly triggered via microbial and damage sensing and acquire an immune specification focused toward the recruitment of neutrophils in the oral mucosa. Moreover, exaggerated activation of stromal cells in disease is linked to excessive recruitment of immune cell populations and destructive inflammatory immune responses.

## Discussion

Herein we present a robust atlas of human oral mucosal tissues in health and in the setting of the common oral inflammatory disease periodontitis. Our work details the complex landscape of oral mucosal cell types and provides comprehensive insights into cell functionality and disease susceptibility at this unique barrier.

A definite strength of our study is the meticulous characterization of patient cohorts with a strict definition of oral health and periodontal disease. Indeed, our healthy volunteer cohort was screened for medical history, medication, tobacco/alcohol/drug use and also evaluated through blood work and detailed oral examination. This robust study design has enabled the creation of an atlas of true systemic and local (oral) health that can serve as a community resource towards understanding cell types and their functionality associated with oral mucosal tissue homeostasis. Our health cohort also serves as a normative baseline against which we can begin to explore changes in cell populations and their transcriptomes in the setting of oral tissue diseases. Furthermore, our evaluation of both tooth-associated (gingiva) and lining (buccal) oral mucosa provides a baseline towards evaluation of diseases associated with these two distinct sites which typically present with unique disease susceptibilities^27,28^.

In the current study, we explored shifts in cell populations and their transcriptomes in the setting of untreated, severe inflammatory disease periodontitis, a disease of the gingival mucosa. Periodontitis is one of the most common human diseases^8^ associated with morbidity and significant global health and economic costs^29^ and is considered an independent risk factor for multiple systemic inflammatory diseases^30^. However, mechanisms underlying disease susceptibility and pathogenesis are not fully understood and no targeted treatment is currently available. Our work here describes an atlas of cell populations, transcriptomes and interactions in this mucosal disease and provides insights into the role of particular cell populations in disease susceptibility and pathogenesis, opening avenues for further mechanistic exploration of this complex disease.

Our findings demonstrate that the gingival mucosa is a particularly active tissue site with unique immunological wiring and enhanced inflammatory responsiveness where stromal cell populations actively contribute to homeostatic inflammation. As such, we not only document increased representation of immune cells at this oral mucosa site (particularly granulocytes/neutrophils) but also enhanced proinflammatory and antimicrobial gene expression by stromal and epithelial cells. Heightened defense activation is consistent with the unique environment of the gingival mucosa, which is heavily exposed to a rich and diverse commensal microbiome^31^ and subject to continuous mechanical stimulation through mastication^32^.

The atlas reveals a previously unappreciated diversity of stromal and epithelial cells at the oral mucosa barrier. We find that several stromal/epithelial populations support prototypic cell functions, including keratinization in the epithelium, matrix accumulation and turnover in fibroblasts and angiogenesis/cell transmigration in endothelial cells. In addition, we document unique stromal cell populations with immune functionality which are particularly expanded in the gingival mucosa. In health, we characterize a gingiva-specific epithelial cell population as well as expanded fibroblast populations expressing inflammatory mediators. Gene expression suggests that these pro-inflammatory populations primarily support antimicrobial responses, recruitment of neutrophils and enhanced tissue turn over. Overall, in health we observe that stromal cells in gingiva are wired towards the induction of inflammatory responses and neutrophil recruitment. Indeed, the gingival mucosa is a site of constant neutrophil transmigration^3^. Importantly, we find that stromal cells become particularly activated in the setting of periodontitis promoting immune cell recruitment and migration into the lesions of disease. In disease, fibroblasts and other stromal cells promote the specific recruitment of neutrophils but also express chemokines known to attract other leukocytes, including lymphocytes. These findings suggest a previously unacknowledged role for the mesenchymal compartment in immune responsiveness in oral homeostasis and disease pathogenesis, similar to recent findings that identify a mesenchymal-inflammatory axis in diseases of other tissue compartments^33,34^. Importantly, an unreported aspect of stromal hyper-responsiveness in gingival mucosal health and disease is the particular specification of gingival stromal cells towards the recruitment of neutrophils.

Our findings also further reinforce the critical role of neutrophils in oral mucosal immunity. We document that many of the immune-related pathways upregulated in stromal and epithelial cells in gingival health and disease are related to neutrophil homing. In fact, some of the top upregulated genes in gingival stromal and epithelial cells include *CSF3, CXCL1, CXCL2*, and *CXCL8* which are specific for neutrophil recruitment. Indeed, at steady state, neutrophils are the immune cell type with significantly higher representation by scRNA seq in oral mucosa compared to all other healthy human barrier tissues investigated. In periodontal disease, we further document significant recruitment of neutrophils in tissue lesions by flow cytometry, which is indeed a well appreciated phenomenon^35,36^. scRNA seq captures the increase in tissue neutrophils in periodontitis, albeit without reaching statistical significance. This indicates a technical limitation in capturing tissue neutrophil responses, likely due to the abrupt drop in transcriptional activity of mature neutrophils after leaving the bone marrow, as well as the sensitivity of neutrophils during tissue preparation and isolation steps^14^. Finally, expression of known disease susceptibility genes within oral tissues reflects a predominant expression of Mendelian-disease associated genes within tissue neutrophils. Indeed, it is well recognized that patients with genetic neutrophil defects commonly develop dominant severe oral phenotypes including aggressive periodontitis in children and recurrent oral ulcerations in lining mucosa^15,37^. Furthermore, our atlas allows us to evaluate the expression of genes related to periodontal susceptibility in common forms of periodontitis, identified through GWAS studies. Expression of susceptibility genes within the tissue of interest reveals cell expression in a variety of cell populations and opens avenues for investigation of diverse pathways of pathogenesis in common forms of disease.

Finally, we utilized our atlas to explore expression of microbial receptors and damage sensing molecules in the oral mucosa. At this barrier, a rich commensal microbiome is present in health which may participate in tissue homeostasis and is decidedly involved in the pathogenesis of common oral mucosal diseases including periodontitis, candidiasis and oral ulcers, suggesting a primary role for host-microbial interaction in oral tissue pathophysiology. Furthermore, as multiple commensals and pathogens are first encountered at the oral mucosa prior to entry into the gastrointestinal and respiratory tracts, understanding initial recognition of microbial elements at this barrier becomes particularly important. In addition, the masticatory mucosa, which includes the gingiva, is continuously exposed to mechanical stress and cell damage through mastication^38^. We therefore created an atlas for expression of major bacterial, fungal and viral receptors and sensing molecules within the cell populations of the oral mucosa. We determine distinct receptor expression at buccal and gingival mucosal sites, potentially reflecting unique disease and infection susceptibilities at each oral barrier. Furthermore, modulation of microbial expression in the setting of periodontitis suggests altered host-microbial interactions in the presence of inflammation. Importantly, the expression of microbial sensing and damage receptors within stromal cells and further upregulation of such receptors in the setting of disease supports the concept that the stromal compartment receives diverse microbial and damage signals to promote homeostatic inflammation, but upon exacerbation causes inflammatory tissue destruction and disease.

Taken together, our oral mucosal tissue atlas provides a comprehensive characterization of cell populations and states in health and periodontitis and provides insights into cell functionality and intercellular interactions in disease pathogenesis. Furthermore, our work reveals unique functionality of stromal cell populations participating in homeostatic immunity and inflammation at this barrier.

## Methods

### Human studies

#### Patient recruitment and ethical approval

All healthy volunteers and patients participating in this study were enrolled in an IRB approved clinical protocol at the NIH Clinical Center (ClinicalTrials.gov #NCT01568697) and provided written informed consent for participation in this study. All participants were deemed systemically healthy based on detailed medical history and select laboratory work up.

Inclusion criteria for this study were: ≥ 18 years of age, a minimum of 20 natural teeth, and in good general health. Exclusion criteria were: history of Hepatitis B or C, history of HIV, prior radiation therapy to the head or neck, active malignancy except localized basal or squamous cell carcinoma of the skin, treatment with systemic chemotherapeutics or radiation therapy within 5 years, pregnant or lactating, diagnosis of diabetes and/or HbA1C level >6%, > 3 hospitalizations in the last 3 years, autoimmune disorder such as Lupus, Rheumatoid Arthritis, etc. Additional exclusion criteria included the use of any of the following in the 3 months before study enrollment: systemic (intravenous, intramuscular, or oral) antibiotics, oral, intravenous, intramuscular, intranasal, or inhaled corticosteroids or other immunosuppressants, cytokine therapy, methotrexate or immunosuppressive chemotherapeutic agents, large doses of commercial probiotics (greater than or equal to 10^8^ colony-forming units or organisms per day), or use of tobacco products (including e-cigarettes) within 1 year of screening. In addition to systemic screening, oral health was assessed and only participants with no soft tissue lesions, no signs/symptoms of oral/dental infection and no/minimal gingival inflammation were considered for the health group. Inclusion criteria for periodontal disease group were moderate to severe periodontitis (>5mm probing depth in more than 4 interproximal sites) and visible signs of tissue inflammation including erythema/edema, and bleeding upon probing.

#### Biopsy collection

During study visits participants received detailed intraoral soft tissue and periodontal examination, which included full mouth probing depth and clinical attachment loss measurements (PD, measures of bone destruction) and bleeding on probing (BOP, measure of mucosal inflammation). Standardized 4mm long x 2mm wide gingival collar biopsies and/or 4mm buccal punch biopsies were obtained under local anesthesia. Health group buccal and gingival biopsies were obtained from individuals that met criteria for oral health and in areas without BOP and with PD <3mm. Biopsies of periodontitis patients were obtained from areas of severe inflammation and bone loss (BOP positive and PD >5mm). Each biopsy was analyzed separately and not pooled. Biopsies were either placed into RPMI medium to be processed into single cell suspensions or fixed with 10% zinc formalin (American MasterTech Scientific) for histology. Donor information can be found in Supplemental Table 1.

### Histology

For histology, tissues were placed in zinc formalin for 18-24 hours and then formalin was replaced with 70% ethanol (Sigma). Formalin-fixed tissues were embedded in paraffin and sectioned into 5mm sections, deparaffinized, and rehydrated, followed by Hematoxylin & Eosin staining (H&E) and immunofluorescence staining. Immunofluorescence was performed via heat-mediated antigen retrieval with a Tris-EDTA buffer (10mM Tris, 1mM EDTA, pH 9.0) in a Retriever 2100 pressure cooker (Electron Microscopy Sciences). Slides were incubated overnight at 4°C with the following primary antibodies: anti-vimentin (abcam, ab24525, 1:100 dilution), anti-CD45 (Cell Signaling, 13917, 1:100), anti-CD31 (Cell Signaling, 3528, 1:250), and anti-keratin 5 (LSBio, LS-C22715, 1:500). Alexa Fluor-conjugated secondary antibodies were used to detect the primary antibodies (ThermoFisher, Alexa 488, 546, 594, & 647, all secondary antibodies at 1:300 dilution), and nuclei stained with DAPI (ThermoFisher, 1µg/mL) prior to covering with ProLong Gold Antifade Mountant (ThermoFisher). Slides were imaged on the Leica TCS SP8 X (Leica) located in the National Institute of Arthritis and Musculoskeletal and Skin Diseases (NIAMS) Light Imaging Core and processed on LAS X software (Leica).

### Biopsy preparation into single cell suspension

#### 10X

To prepare single cell suspensions biopsies were minced and digested using Collagenase II (Worthington Biochemical Corporation) and DNase (Sigma) and processed through the gentleMACS Dissociator (Miltenyl) utilizing a nasal mucosa protocol for processing^39^. Following tissue dissociation, cells were passed through a 70μm filter (Falcon, Corning), washed, and counted with a Cellometer Auto 2000 (Nexcelom).

#### Flow cytometry

Human gingival tissues were minced and digested for 50 minutes at 37°C with Collagenase IV (Gibco) and DNAse (Sigma). A single-cell suspension was then generated by mashing digested samples through a 70μm filter (Falcon, Corning), as previously described^35^.

### Flow cytometry of human oral mucosal tissue samples

Single-cell suspensions from gingival tissues were incubated with mouse serum (Jackson ImmunoResearch Lab) and fluorochrome-conjugated antibodies against surface markers in PBS with 2.5% FBS, for 20 minutes at 4°C in the dark, and then washed. Dead cells were excluded with Live/Dead fixable dye (Amcyan, 1:100, Invitrogen). Anti-human antibody information used for staining is described in Supplemental Table 9. Cell acquisition was performed on a BD LSRFortessa machine using FACSDiVa software (BD Biosciences). Data were analyzed with FlowJo software (TreeStar).

### 10X Sequencing

Single-cell suspensions were loaded onto a 10X Chromium Controller (10X Genomics) and library preparation was performed according to the manufacturer’s instructions for the 10X Chromium Next GEM Single Cell Library kit v3 (10X Genomics). Libraries were then pooled in groups of 4 and sequenced on 4 lanes on a NextSeq500 sequencer (Illumina) using 10X Genomics recommended reads configuration.

### 10X Data alignment

Read processing was performed using the 10X Genomics workflow. Briefly, the Cell Ranger Single-Cell Software Suite v3.0.1 was used for demultiplexing, barcode assignment, and unique molecular identifier (UMI) quantification (http://software.10xgenomics.com/single-cell/overview/welcome). The reads were aligned to the hg38 human reference genome (Genome Reference Consortium Human Build 38) using a pre-built annotation package obtained from the 10X Genomics website (https://support.10xgenomics.com/single-cell-gene-expression/software/pipelines/latest/advanced/references). All 4 lanes per sample were merged using the ‘cellranger mkfastq’ function and processed using the ‘cellranger count’ function.

### 10X Data import

For oral datasets, filtered feature barcode matrices generated by the cellranger pipeline were used as input into a Seurat-compatible format using the function ‘Read10X’ and ‘CreateSeuratObject’.

For previously published skin^11^, lung^13^ and ileum^12^ datasets, files downloaded from GEO Accession Viewer were extracted and imported into a Seurat-compatible object. All quality control, normalization, and downstream analysis was performed using the R package Seurat^40^ (ver. 3.2.2, https://github.com/satijalab/seurat) unless otherwise noted.

### 10X Data quality control, normalization and integration

Cells were filtered to remove any that expressed fewer than 200 genes, more than 5,000 genes and those with more than 15% mitochondrial gene expression. Normalization steps were performed using the Seurat function ‘NormalizeData’. A total of three unique dataset integrations were performed for data analysis using the Seurat functions ‘FindIntegrationAnchors’ and ‘IntegrateData’: 1. buccal and gingival health, 2. gingival health and disease, and 3. oral (buccal and gingival health), skin, lung, and ileum. For the third group of samples described above, all cohorts were randomly downsampled using the base R function ‘sample’ such that each group (oral, skin, lung, ileum) contained the same number of cells for global comparison.

### 10X Dimensionality reduction, cell clustering and identification

In order to select for the principal component (PC) that explain the most variance, variance contributed by each PC was visualized using an elbow plot. Informative PCs were then used for downstream analyses. Louvain clustering was performed with the ‘FindClusters’ function using the first 50 PCs. Differential gene expression was used to guide manual annotation of clusters (‘FindAllMarkers’ function). The transcriptomic signature of immune cells was further investigated using the Monaco reference transcriptomic database^41^ of pure cell types via SingleR^42^ (ver. 1.0.6, https://github.com/dviraran/SingleR). To eliminate overlap of individual cells which may obfuscate gene expression patterns, UMAPs for select cell subtypes were tessellated by a regular grid of hexagons, the number of cells within each hexagon were counted, and the average expression of cells within each hexagon was represented by a color ramp via schex (ver. 1.0.55, https://github.com/SaskiaFreytag/schex).

### 10X Pathway analysis and receptor-ligand signaling inference

Gene set over-representation analysis of significantly upregulated cluster-defining genes was performed using the gsfisher R package (ver. 0.2, https://github.com/sansomlab/gsfisher/), and filtered to include only ‘biological process’ gene sets obtained from the GO database.

To predict cell-cell interactions using expression data, we utilized NicheNet^43^ (ver. 1.0.0, https://github.com/saeyslab/nichenetr) which combined gene expression data of cells from our sequencing cohort with a database of prior knowledge on signaling and gene regulatory networks. The integrated Seurat object containing healthy and diseased gingival tissue was used as input into the NicheNet Seurat wrapper (‘nichenet_seuratobj_aggregate’).

### Statistics

To assess significance of proportion changes across tissue types and flow cytometry results, if both sample groups passed the Shapiro-Wilk normality test, an unpaired t-test with Welch’s correction was used. If one or more groups did not pass the normality test, a Mann-Whitney test was used. When assessing differentially expressed genes with Seurat, a non-parametric Wilcoxon rank sum test was used.

## Acknowledgements

This study was funded in part by the intramural programs of NIH/NIDCR (NMM) and NIH/NIAMS (MM), by extramural grants NIH/NIDCR DE025046 (KD), DE029436 and DE028561 (GH), FONDECYT #11180389 through the National Agency of Research and Development (ANID), Chile (ND), and a PhD studentship from the Barbour Foundation (SW). MH is funded by Wellcome (WT107931/Z/15/Z), The Lister Institute for Preventive Medicine and Newcastle NIHR Biomedical Research Centre (BRC). This work was also made possible through use of the Genomics and Computational Biology Core (NIDCR, ZIC DC000086) and Combined Technical Research Core (NIDCR, ZIC DE000729-09). Figure illustrations were created in part with Biorender.com and by Alan Hoofring (Lead Medical Illustrator, NIH Medical Arts). The authors also thank Dr. Tassos Sfondouris for clinical support.

## Author Contributions

DWW and NMM conceived of the study and wrote the manuscript. DWW, TGW, LB, ND were involved in sample acquisition and preparation and flow cytometry experiments and analysis. DWW performed analysis of 10x data and prepared all figures. OA, APS performed histological staining. SW, DM, MH, KD, GH consulted on experiments and data analysis and critically reviewed and edited the manuscript. NMM supervised the study.

## Data availability

Raw and processed single-cell RNA sequencing datasets have been deposited in the NCBI GEO database and can be accessed via the following link; GSE164241, https://www.ncbi.nlm.nih.gov/geo/query/acc.cgi?acc=GSE164241)

## Supplemental Figure Legends

**Supplemental Figure 1 (related to Figure 1). a**. Violin plots showing the number of features, RNA counts and percent mitochondrial transcripts found in gingival health (GM), buccal health (BM), and periodontal disease (PD) patient samples prior to quality control. **b**. Violin plots showing the number of features, RNA counts and percent mitochondrial transcripts following quality control. **c**. Bar graphs showing the number of cells that were originally sequenced (preQC) and the number of cells included for analysis (postQC) for each patient sample. **d**. Heatmap showing the average Z-scored expression of the top differentially regulated genes for the major cell types identified in healthy gingival and buccal tissues.

**Supplemental Figure 2 (related to Figure 2). a**. UMAP depicting the number of clusters identified in healthy gingival and buccal tissue by the Seurat clustering algorithm with a resolution of 1. Clusters 17 and 30 were contaminated with red blood cells and were removed from the dataset prior to analyses. **b-d**. Heatmaps depicting average gene expression for each gene associated with the GO term on the right that was enriched in each individual endothelial **(b)**, epithelial **(c)** and fibroblast **(d)** cluster.

**Supplemental Figure 3 (related to Figure 3). a**. Example gating strategy for flow cytometry analysis in Figure 3h. **b**. Proportion plots showing the percentage of each immune cell type in each barrier tissue. **p<0.01 based on a one-way ANOVA and Dunnett’s multiple comparisons test

**Supplemental Figure 4 (related to Figure 4,5). a**. UMAP depicting the number of clusters identified in healthy and disease gingival tissue by the Seurat clustering algorithm with a resolution of 1. Clusters 17, 24 and 29 were contaminated with red blood cells and were removed from the dataset prior to analyses. **b-c**. Immunohistochemistry of healthy **(b)** and periodontitis **(c)** tissues stained with myeloperoxidase, an enzyme abundantly expressed by neutrophils. Scale bar: 100μm.

**Supplemental Figure 5 (related to Figure 6). a-c**. Dot plots depicting the expression of cluster-defining genes and percentage of cells expressing each gene for endothelial **(a)**, epithelial **(b)**, and fibroblast **(c)** populations, respectively. Expression values are Z-scored averages.

**Supplemental Figure 6 (related to Figure 6). a**. Dot plot showing tissue- and cell-type specific expression of top ligands identified by NicheNet. **b**. Dot plot showing periodontitis immune cell expression of receptors identified by NicheNet. **c**. Heatmap of prior interaction potential between receptors and ligands identified by NicheNet and used to generate alluvial plots in Figure 6g. **d**. Heatmap of inferred downstream genes that are regulated by ligand binding, estimated by the prior model in NicheNet.

**Supplemental Figure 7 (related to Figure 6). a**. Dot plots of chemokine receptor expression in immune cells in health and disease (left) and expression of chemokine ligand expression in general cell types in health and disease (right). Colored lines extending from receptors to ligands represent known interactions. Expression of chemokine receptors was simplified in Figure 6h based on this dot plot.

**Supplemental Figure 8 (related to Figure 7). a-b**. Dot plots showing expression of TLRs (green), NLRs (brown), CLR/RLRs (orange), DAMPs (blue), DAMP receptors (cyan), SARS-CoV-2 viral entry factors (red), and viral entry factors for viruses that lead to oral pathology (purple) in major cell types of gingival and buccal health **(a)** and in gingival health and disease **(b)**.

## Supplemental Tables

**Table S1:** Demographic information of all patients analyzed with single cell RNA sequencing

**Table S2:** List of cluster defining genes for endothelial, epithelial and fibroblasts in Figure 2

**Table S3:** List of cluster defining genes for major immune cell types identified in Figure 3

**Table S4:** Demographic information of all patients analyzed with flow cytometry

**Table S5:** List of gene ontology pathway information described in Figure 6

**Table S6:** Cell and source information for comparative single cell sequencing analysis of barrier tissues

**Table S7:** Wilcoxon rank sum test results for expression of PRRs, DAMPs, DAMP-Rs and viral entry factors. Comparison between healthy gingival and buccal tissues.

**Table S8:** Wilcoxon rank sum test results for expression of PRRs, DAMPs, DAMP-Rs and viral entry factors. Comparison between healthy and diseased gingival tissues.

**Table S9:** Antibody list

