## Supplemental Figures for "Single-cell atlas of human oral mucosa reveals a stromal-neutrophil axis regulating tissue immunity in health and inflammatory disease"

### Supplemental Figure 1

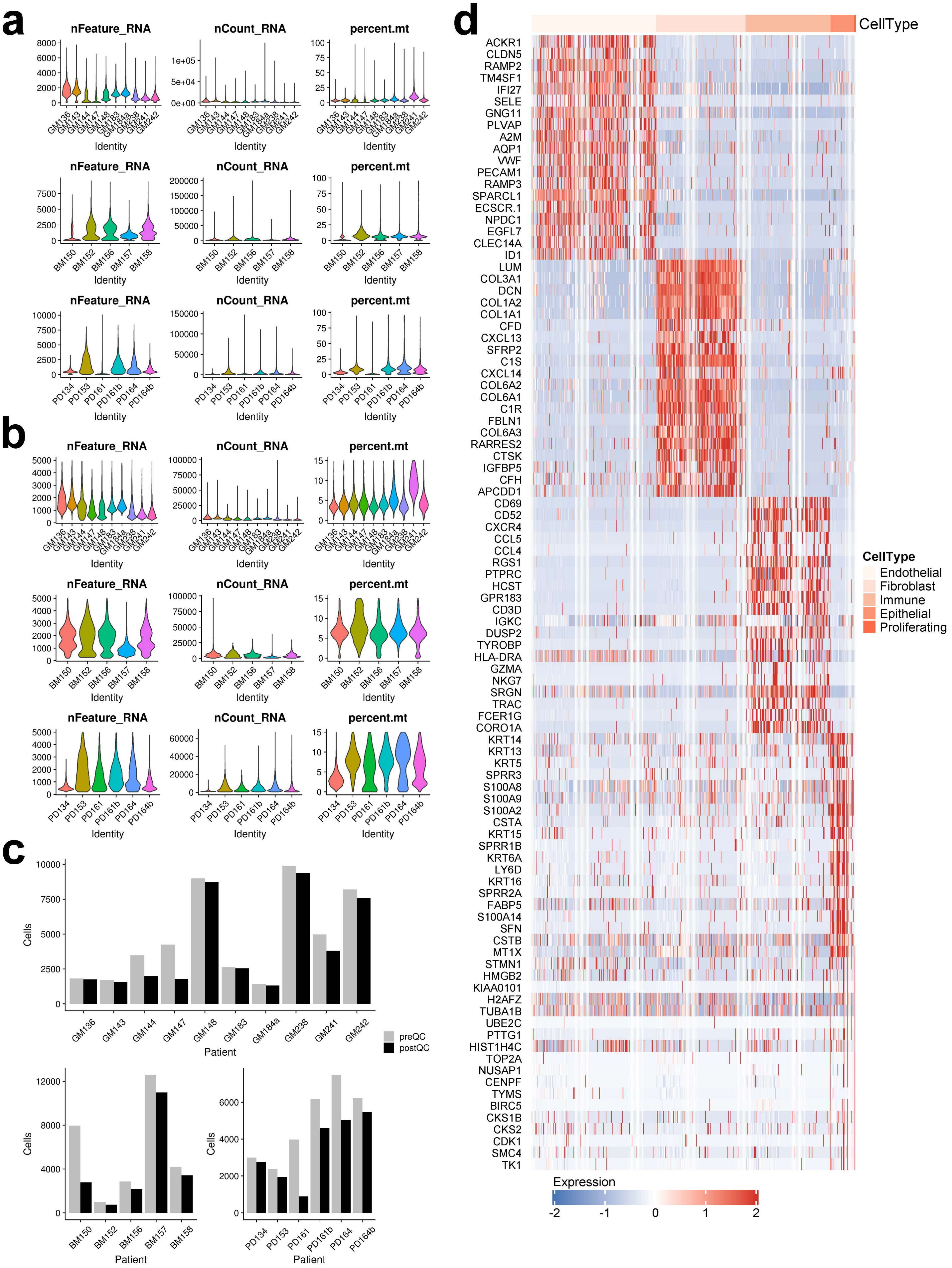

### Supplemental Figure 2

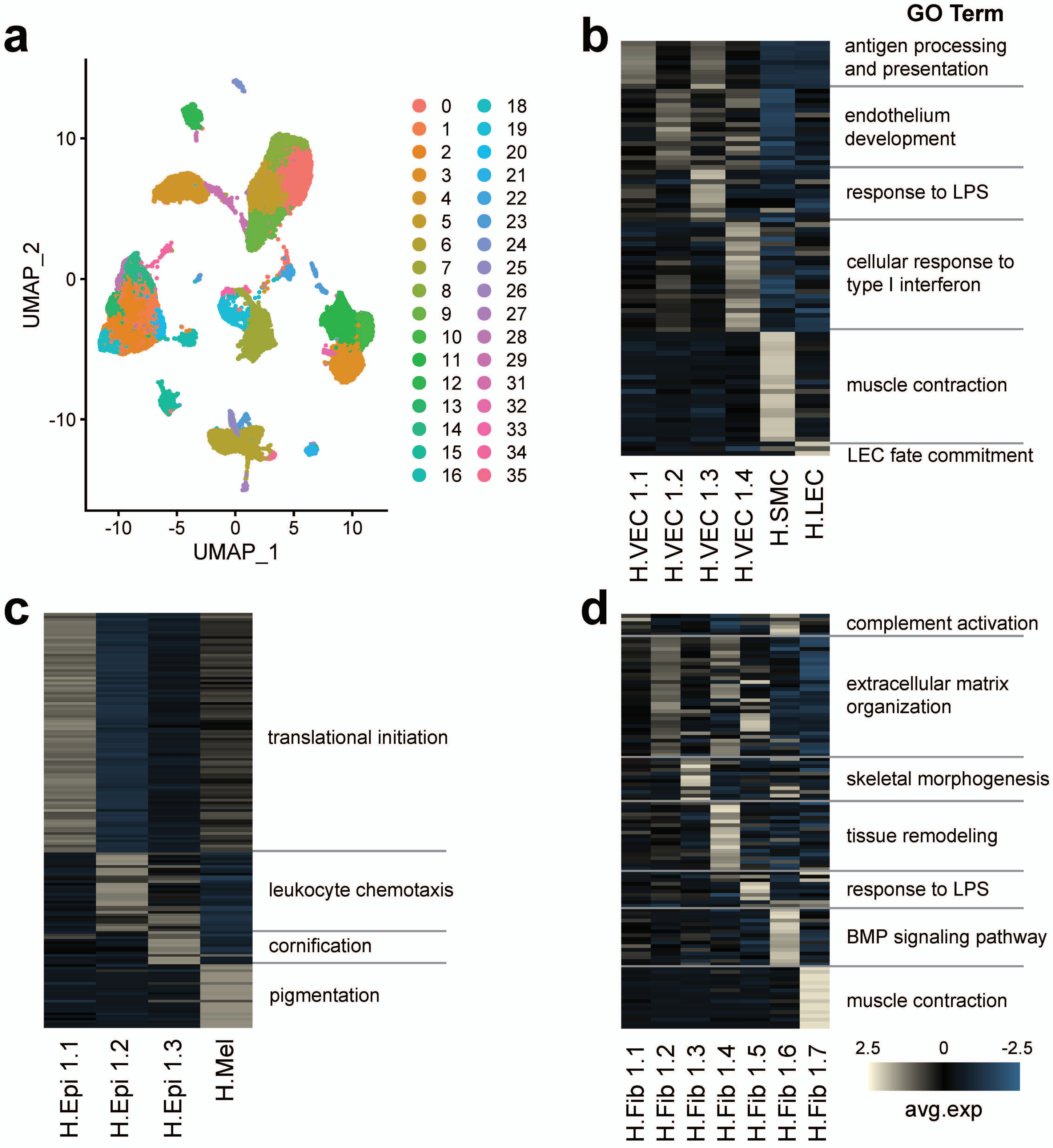

### Supplemental Figure 3

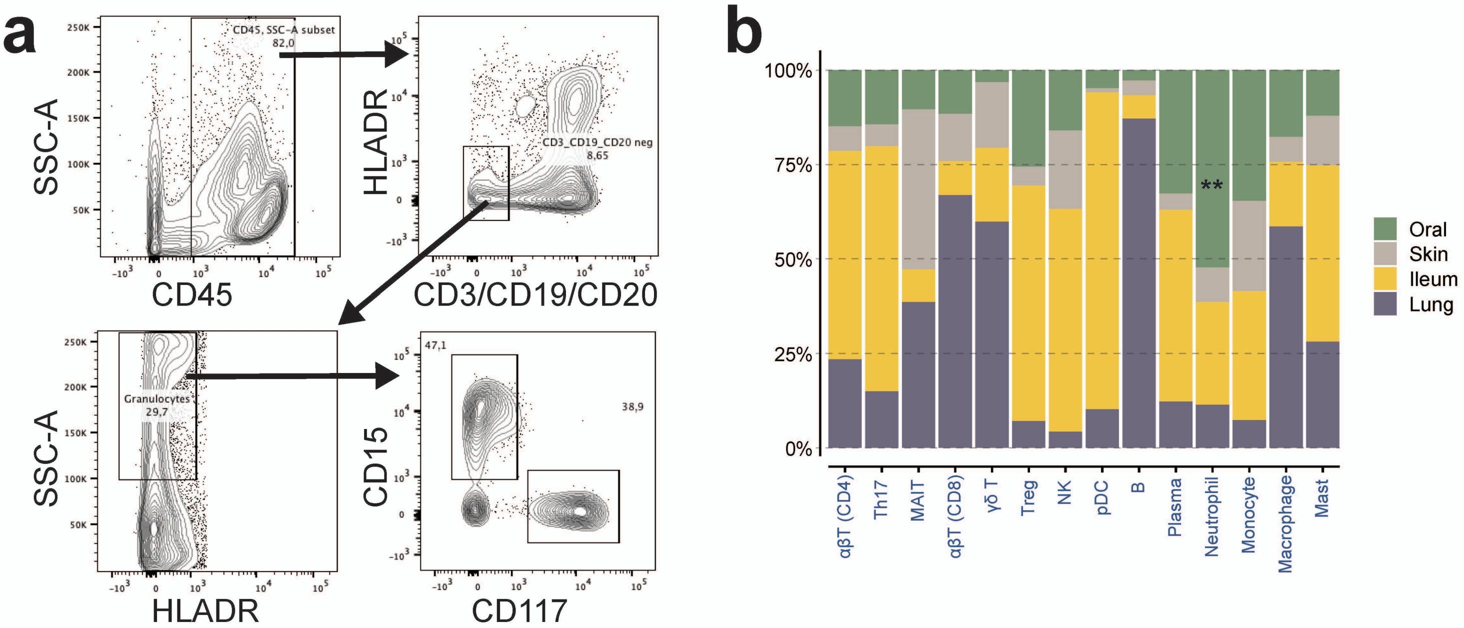

Supplemental Figure 4

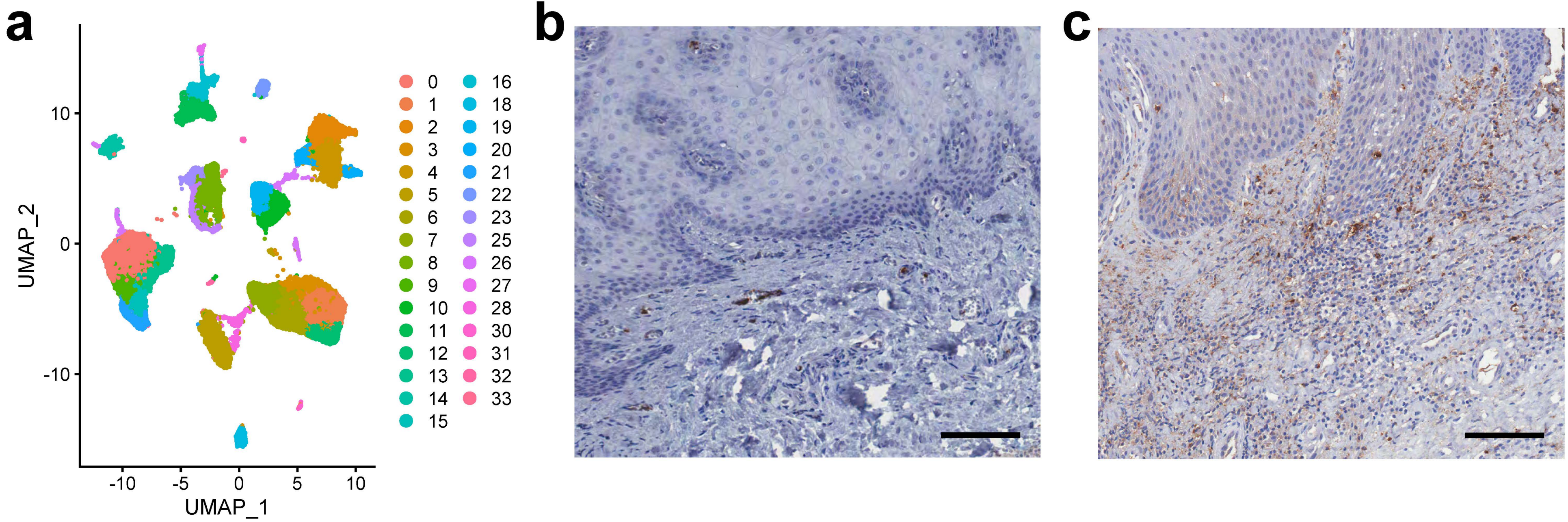

### Supplemental Figure 5

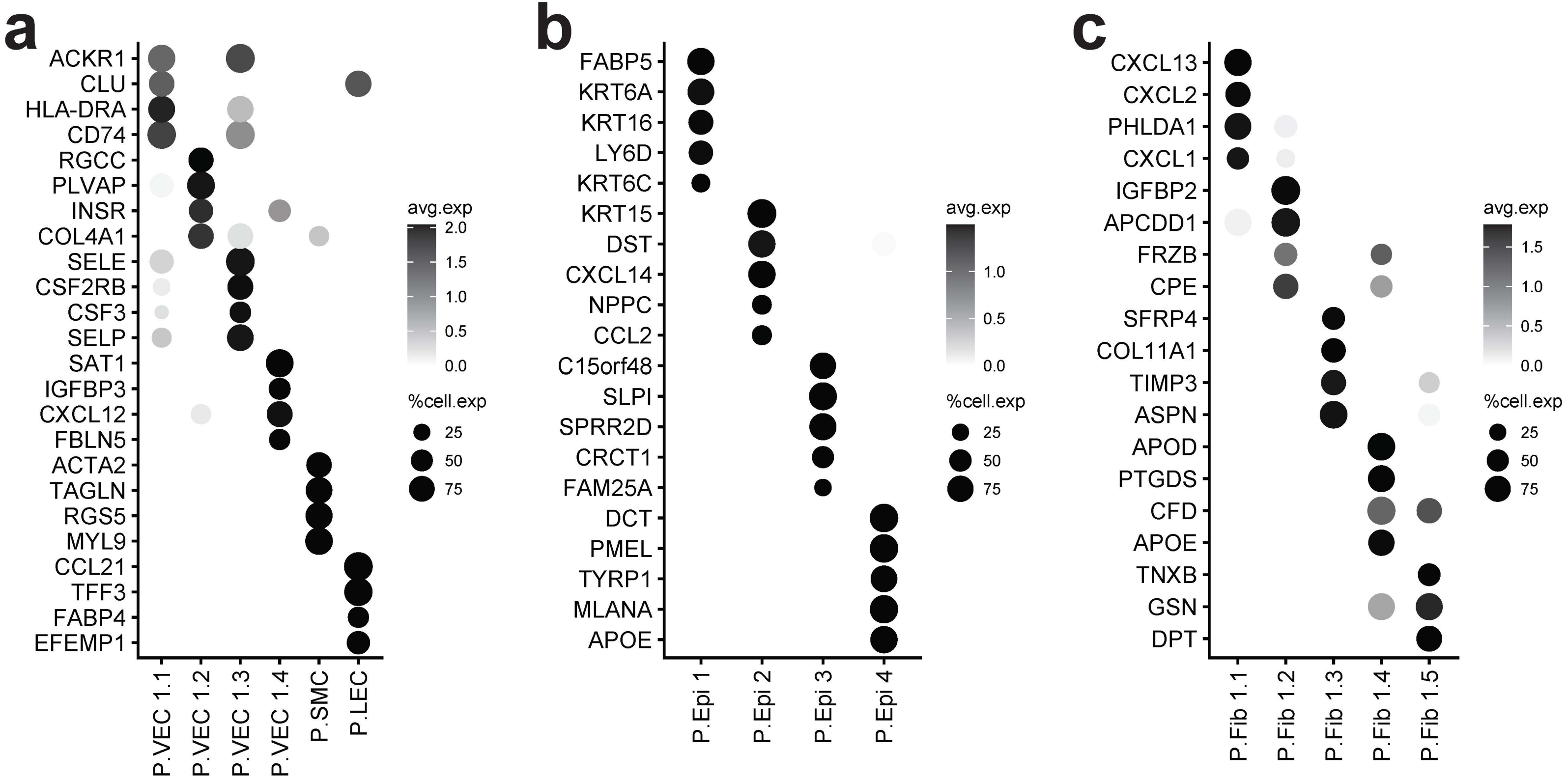

### Supplemental Figure 6

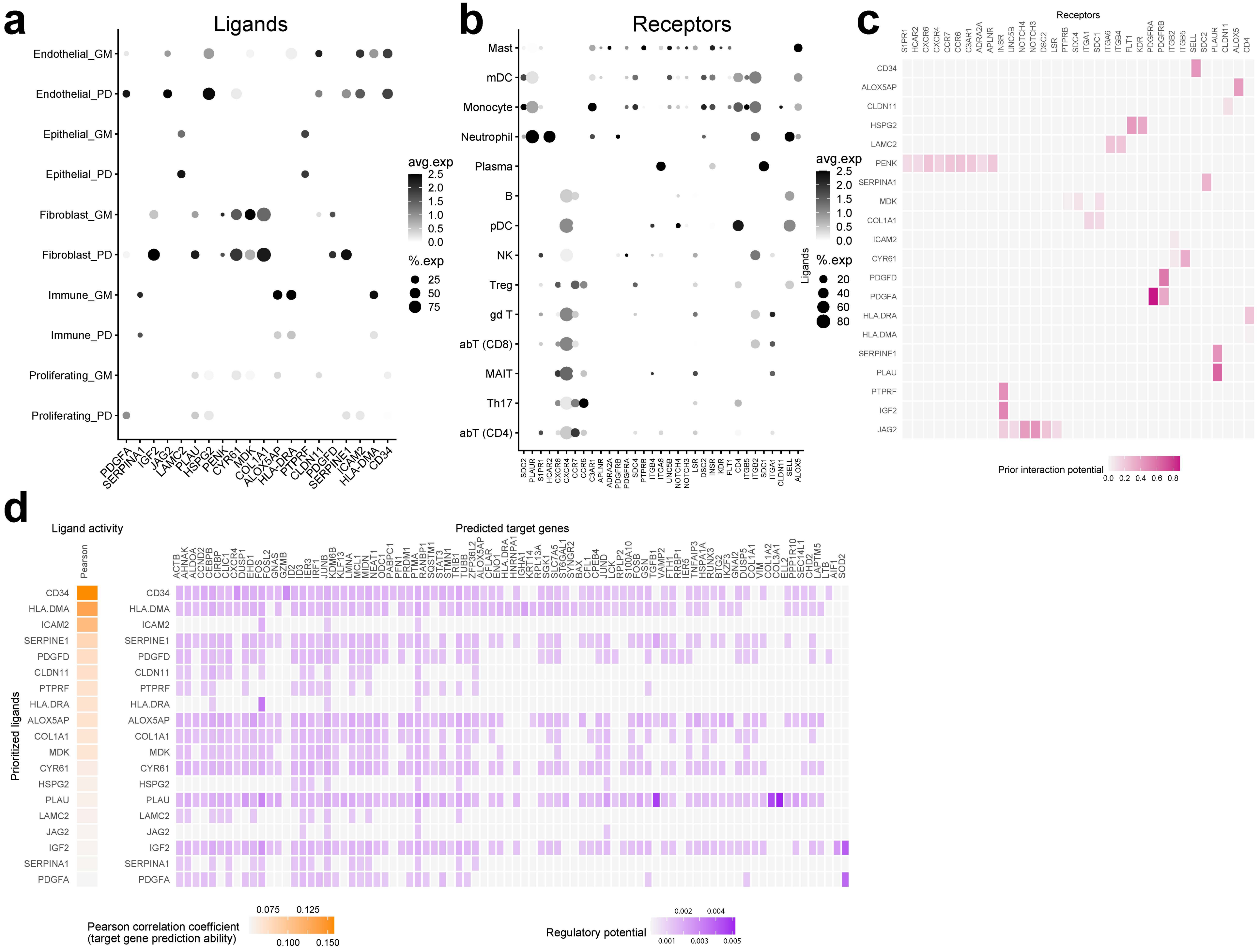

### Supplemental Figure 7

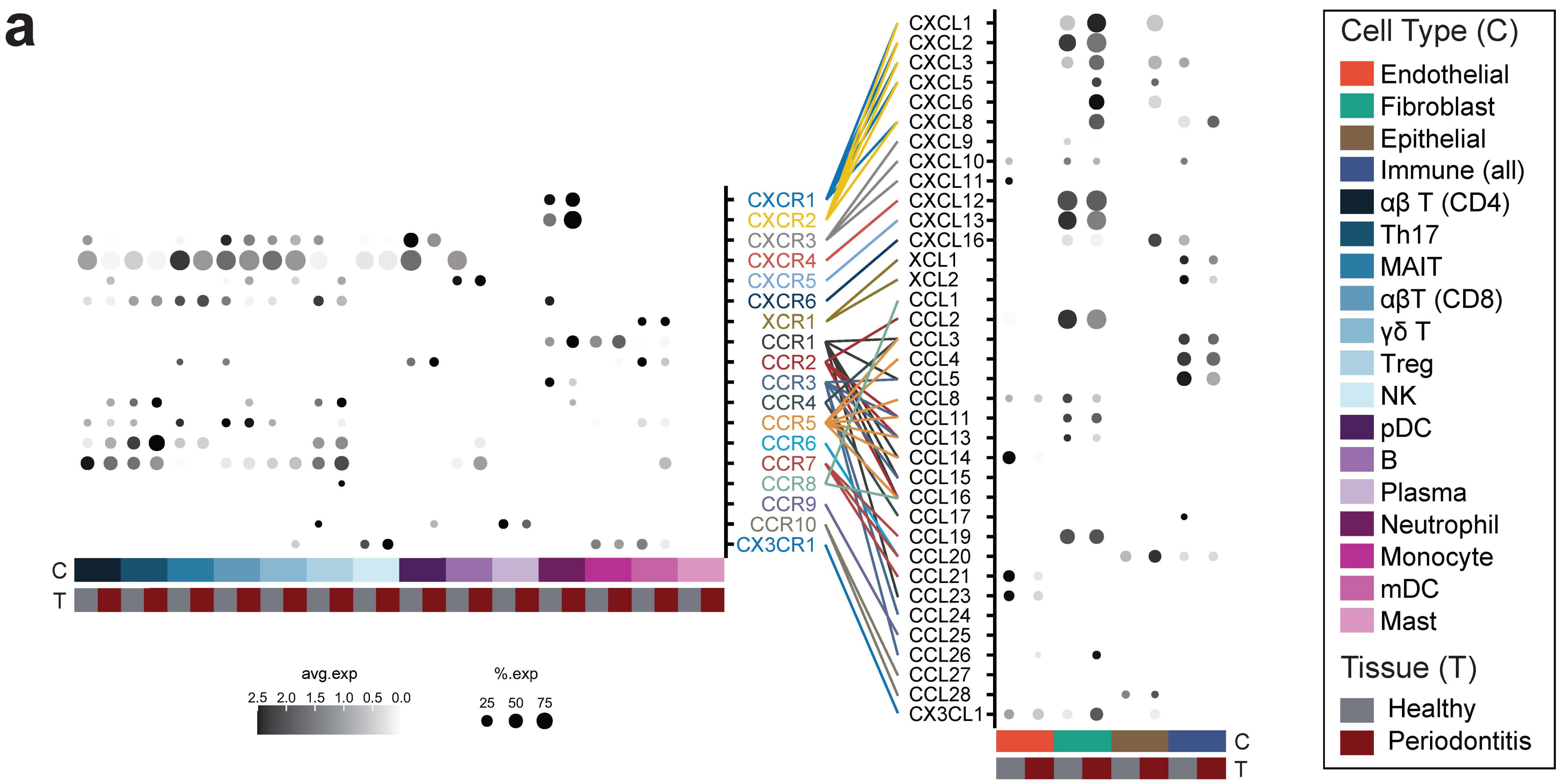

### Supplemental Figure 8

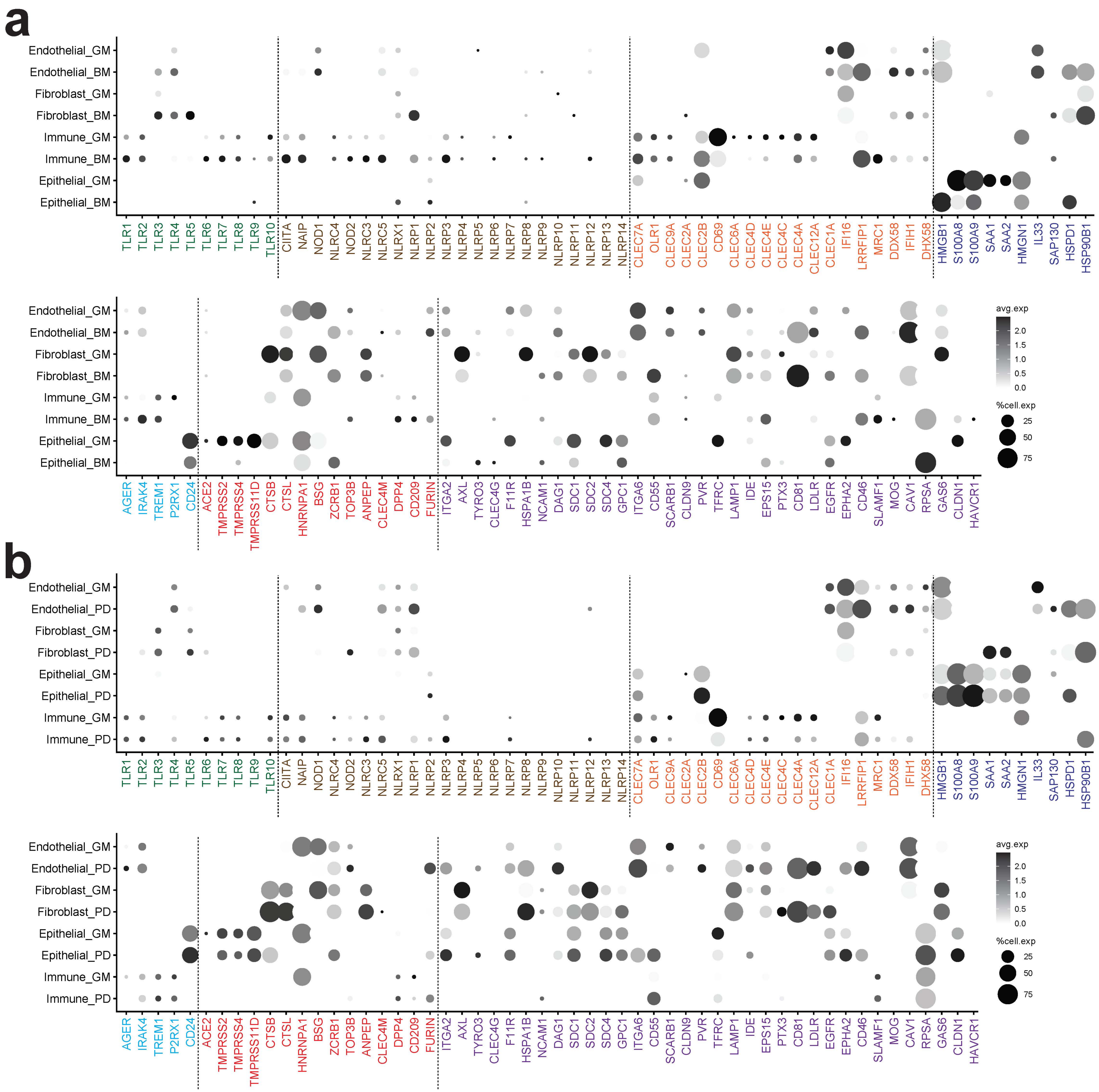
